# SARS-CoV-2 variants B.1.351 and B.1.1.248: Escape from therapeutic antibodies and antibodies induced by infection and vaccination

**DOI:** 10.1101/2021.02.11.430787

**Authors:** Markus Hoffmann, Prerna Arora, Rüdiger Groß, Alina Seidel, Bojan Hörnich, Alexander Hahn, Nadine Krüger, Luise Graichen, Heike Hofmann-Winkler, Amy Kempf, Martin Sebastian Winkler, Sebastian Schulz, Hans-Martin Jäck, Bernd Jahrsdörfer, Hubert Schrezenmeier, Martin Müller, Alexander Kleger, Jan Münch, Stefan Pöhlmann

## Abstract

The global spread of SARS-CoV-2/COVID-19 is devastating health systems and economies worldwide. Recombinant or vaccine-induced neutralizing antibodies are used to combat the COVID-19 pandemic. However, recently emerged SARS-CoV-2 variants B.1.1.7 (UK), B.1.351 (South Africa) and B.1.1.248 (Brazil) harbor mutations in the viral spike (S) protein that may alter virus-host cell interactions and confer resistance to inhibitors and antibodies. Here, using pseudoparticles, we show that entry of UK, South Africa and Brazil variant into human cells is susceptible to blockade by entry inhibitors. In contrast, entry of the South Africa and Brazil variant was partially (Casirivimab) or fully (Bamlanivimab) resistant to antibodies used for COVID-19 treatment and was less efficiently inhibited by serum/plasma from convalescent or BNT162b2 vaccinated individuals. These results suggest that SARS-CoV-2 may escape antibody responses, which has important implications for efforts to contain the pandemic.

## INTRODUCTION

The pandemic spread of severe acute respiratory syndrome coronavirus 2 (SARS-CoV-2), the causative agent of coronavirus disease 2019 (COVID-19), is ravaging economies and health system worldwide and has caused more than 2.3 million deaths ((WHO), 2020). The identification of antivirals by drug repurposing was so far largely unsuccessful. Remdesivir, an inhibitor of the viral polymerase, is the only antiviral with proven efficacy (Beigel et al., 2020). However, the clinical benefit reported for Remdesivir treatment is moderate and has been called into question (Consortium et al., 2020; Wang et al., 2020). Recombinant antibodies, which target the viral spike protein (S) and neutralize infection in cell culture and animal models (Baum et al., 2020a; Chen et al., 2020), have been granted emergency use authorization (EUA) and may provide a valuable treatment option in the absence of other antivirals. In contrast to the moderate success in the area of antivirals, protective mRNA- and vector-based vaccines encoding the SARS-CoV-2 S protein have been approved for human use and are considered key to the containment of COVID-19 (Baden et al., 2021; Polack et al., 2020).

SARS-CoV-2, an enveloped, positive-strand RNA virus that uses its envelope protein spike (S) to enter target cells. Entry depends on S protein binding to the cellular receptor ACE2 and S protein priming by the cellular serine protease TMPRSS2 (Hoffmann et al., 2020b; Zhou et al., 2020) and these processes can be disrupted by soluble ACE2 and serine protease inhibitors (Hoffmann et al., 2020b; Monteil et al., 2020; Zhou et al., 2020). Further, the S protein of SARS-CoV-2 and other coronaviruses is a major determinant of viral cell and species tropism and the main target for the neutralizing antibody response. The genetic information of SARS-CoV-2 has remained relatively stable after the detection of first cases in Wuhan, China, in the winter season of 2019. The only exception was a D614G change in the viral S protein that became dominant early in the pandemic and that has been associated with increased transmissibility (Korber et al., 2020; Plante et al., 2020; Volz et al., 2021). In contrast, D614G has only a moderate impact on SARS-CoV-2 neutralization by sera from COVID-19 patients and by sera from vaccinated individuals (Korber et al., 2020; Weissman et al., 2021).

In recent weeks several SARS-CoV-2 variants emerged that seem to exhibit increased transmissibility and that harbor mutations in the S protein. The SARS-CoV-2 variant B.1.1.7 (UK variant), also termed variant of concern (VOC) 202012/01 or 20I/501Y.V1, emerged in the United Kingdom and was associated with a surge of COVID-19 cases (Leung et al., 2021). Subsequently, spread of the UK variant in other countries was reported (Claro et al., 2021; Galloway et al., 2021). It harbors nine mutations in the S protein, six of which are located in the surface unit, S1, and three are found in the transmembrane unit, S2 (Fig. 1). Exchange N501Y is located in the receptor binding domain (RBD), a domain within S1 that interacts with ACE2, and its presence was linked to increased human-human transmissibility (Leung et al., 2021; Zhao et al., 2021). Variants B.1.351 (20H/501Y.V2, also termed South Africa variant) and B.1.1.248 (P.1., also termed Brazil variant) were also purported to be more transmissible and these variants harbor nine and eleven mutations in their S proteins, respectively, including three changes in the RBD, K417N/T, E484K and N501Y (Fig. 1) (CDC, 2021). These mutations, as well as the N501Y change present in the S protein of the UK variant, may alter host cell interactions and susceptibility to experimental entry inhibitors and antibody-mediated neutralization. However, no functional characterization of the S proteins of UK, South Africa and Brazil variant have been reported in the peer-reviewed literature, with the exception of one study showing that the UK variant exhibits reduced susceptibility to neutralization by sera from COVID-19 patients and vaccinated individuals (Muik et al., 2021) and another study showing that mutations E484K and N501Y, which are both present in the South Africa and Brazil variants, have little effect on neutralization by sera from individuals who were immunized twice with BNT162b2 (Xie et al., 2021).

**Figure 1.**
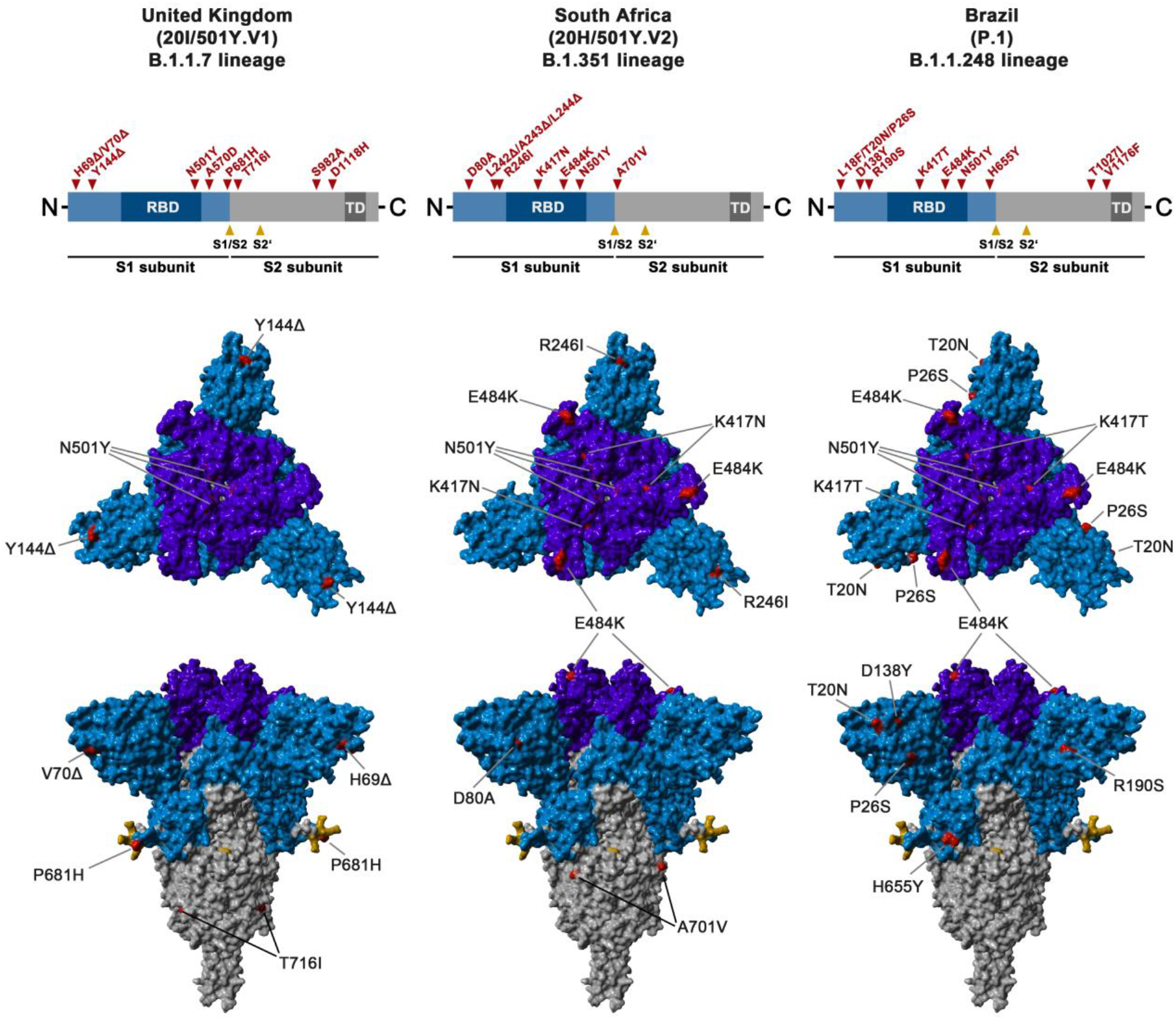
Schematic overview of the S proteins from the SARS-CoV-2 variants under study. The location of the mutations in the context of spike protein domain organization is shown in the upper panel. RBD = receptor binding domain, TD = transmembrane domain. The location of the mutations in the context of the trimer spike protein domain is shown lower panel. Color code: light blue = S1 subunit with RBD in dark blue, grey = S2 subunit, orange = S1/S2 and S2’ cleavage sites, red = mutated amino acid residues.

Here, we show that the S protein of the UK, South Africa and Brazil variants mediate robust entry into human cell lines and that entry is blocked by soluble ACE2 (sACE2), protease inhibitors active against TMPRSS2 and membrane fusion inhibitors. In contrast, monoclonal antibodies with EUA for COVID-19 treatment partially or completely failed to inhibit entry driven by the S proteins of the South Africa and Brazil variants. Similarly, these variants were less efficiently inhibited by convalescent plasma and sera from individuals vaccinated with BNT162b2. Our results suggest that SARS-CoV-2 can evade inhibition by neutralizing antibodies.

## RESULTS

### The spike proteins of the SARS-CoV-2 variants mediate robust entry into human cell lines

We first investigated whether the S proteins of SARS-CoV-2 WT (Wuhan-1 isolate with D614G exchange), UK, South Africa and Brazil variants (Fig. 1) mediated entry into human and non-human primate (NHP) cell lines with comparable efficiency. For this, we used a vesicular stomatitis virus (VSV)-based vector pseudotyped with the respective S proteins. This system faithfully mimics key aspects of SARS-CoV-2 entry into cells, including ACE2 engagement, priming of the S protein by TMPRSS2 and antibody-mediated neutralization (Hoffmann et al., 2020b). The following cell lines are frequently used for SARS-CoV-2 research and were employed as target cells in our study: The African green monkey kidney cell line Vero, Vero cells engineered to express TMPRSS2, the human embryonic kidney cell line 293T, 293T cells engineered to express ACE2, the human lung cell line Calu-3 and the human colon cell line Caco-2. All cell lines tested express endogenous ACE2. In addition, Calu-3 and Caco-2 cells express endogenous TMPRSS2 (Bottcher-Friebertshauser et al., 2011; Kleine-Weber et al., 2018).

All S proteins studied were robustly expressed and mediated formation of syncytia in transfected cells (Fig. 2A). Entry into all cell lines was readily detectable but the relative entry efficiency varied. Particles bearing the S proteins of the SARS-CoV-2 variants entered 293T (Brazil variant) and 293T-ACE2 (South Africa and Brazil variants) cells with slightly reduced efficiency as compared to particles bearing WT S protein, while the reverse observation was made for Calu-3 cells (UK variant). For the remaining cell lines, no significant differences in entry efficiency were observed between SARS-CoV S WT and S proteins from SARS-CoV-2 variants (Fig. 2B). Collectively, these results indicate that the mutations present in the S proteins of UK, South Africa and Brazil variant are compatible with robust entry into human cells.

**Figure 2.**
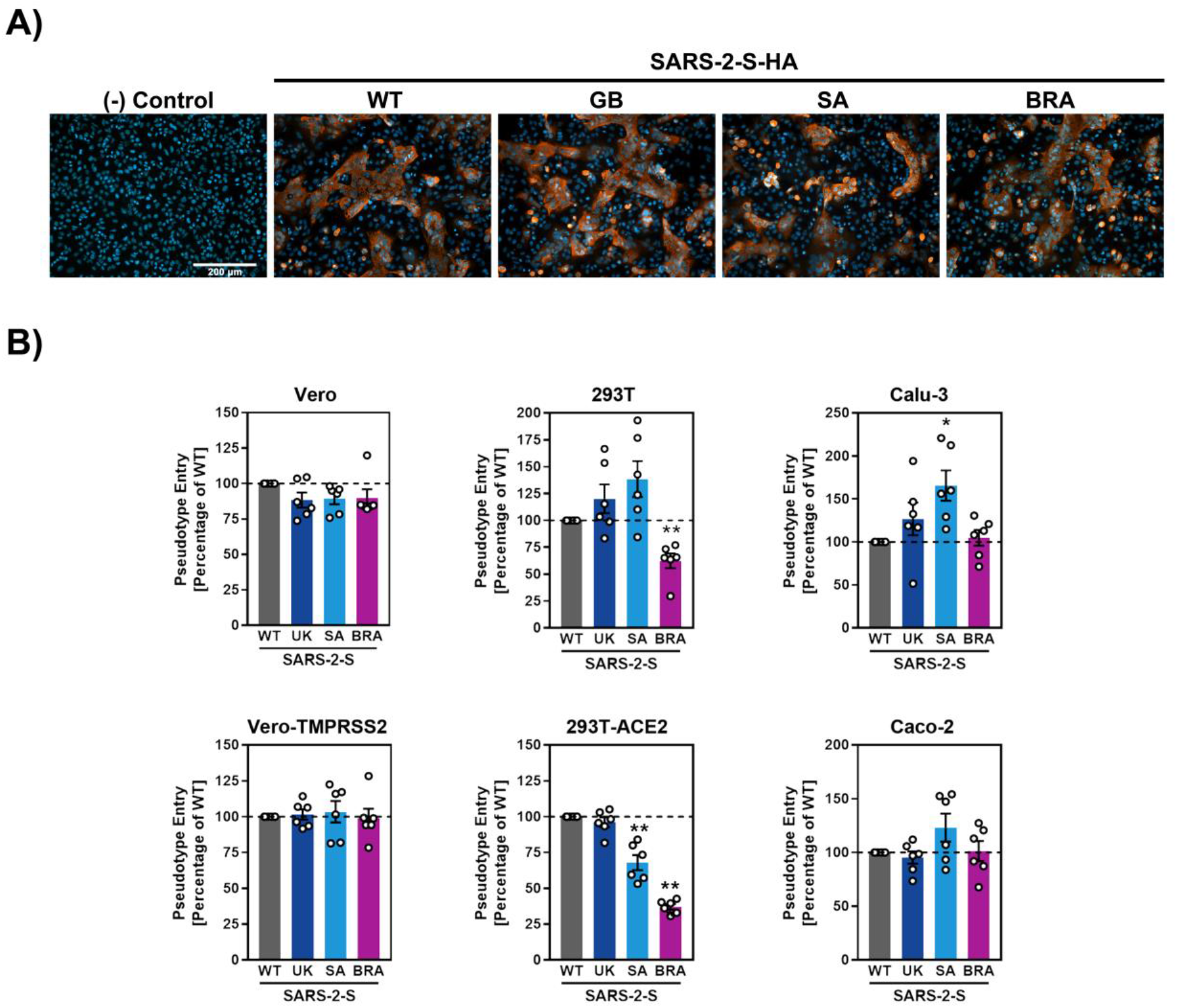
S proteins from SARS-CoV-2 variants drive entry into human cell lines. (A) Directed expression of SARS-CoV-2 S proteins (SARS-2-S) in A549-ACE2 cells leads to the formation of syncytia. S protein expression was detected by immunostaining using an antibody directed against a C-terminal HA-epitope tag. Presented are the data from one representative experiment. Similar results were obtained in four biological replicates. (B) The S proteins of the SARS-CoV-2 variants mediate robust entry into cell lines. The indicated cell lines were inoculated with pseudotyped particles bearing the S proteins of the indicated SARS-CoV-2 variants. Transduction efficiency was quantified by measuring virus-encoded luciferase activity in cell lysates at 16-20 h post transduction. Presented are the average (mean) data from six biological replicates (each conducted with technical quadruplicates). Error bars indicate the standard error of the mean (SEM). Statistical significance was analyzed by one-way analysis of variance (ANOVA) with Dunnett’s posttest. WT = wildtype, GB = Great Britain, SA = South Africa, BRA = Brazil

### The spike proteins of the SARS-CoV-2 variants mediate fusion of human cells

The S protein of SARS-CoV-2 drives cell-cell fusion resulting in the formation of syncytia and this process might contribute to viral pathogenesis (Buchrieser et al., 2021). We employed a cell-cell fusion assay to determine whether the S proteins of UK, South Africa and Brazil variant drive fusion with human cells. For this, the S proteins under study were expressed in effector 293T cells, which were subsequently mixed with target 293T cells engineered to express ACE2 or ACE2 in conjunction with TMPRSS2. The S protein of SARS-CoV was included as control. The SARS-CoV S protein failed to mediate fusion with target cells expressing ACE2 only but efficiently drove fusion with cells expressing ACE2 and TMPRSS2 (Fig. 3A). Similar results were obtained by microscopic examination of A549-ACE2 and A549-ACE2/TMPRSS2 cells transfected to express the respective S proteins (Fig. 3B). These findings are in agreement with the documented requirement for an exogenous protease for SARS-CoV S driven cell-cell fusion under the experimental conditions chosen (Hoffmann et al., 2020a). In contrast, the SARS-CoV-2 S protein mediated efficient membrane fusion in the absence of TMPRSS2 expression in target cells (Fig. 3A,B) and this property is known to depend on the multibasic S1/S2 site of this S protein which is absent in SARS-CoV S (Hoffmann et al., 2020a). Finally, the S proteins of all SARS-CoV-2 variants tested facilitated cell-cell fusion with similar (UK) or slightly reduced (South Africa, Brazil) efficiency as compared to WT S protein (Fig. 3A,B).

**Figure 3.**
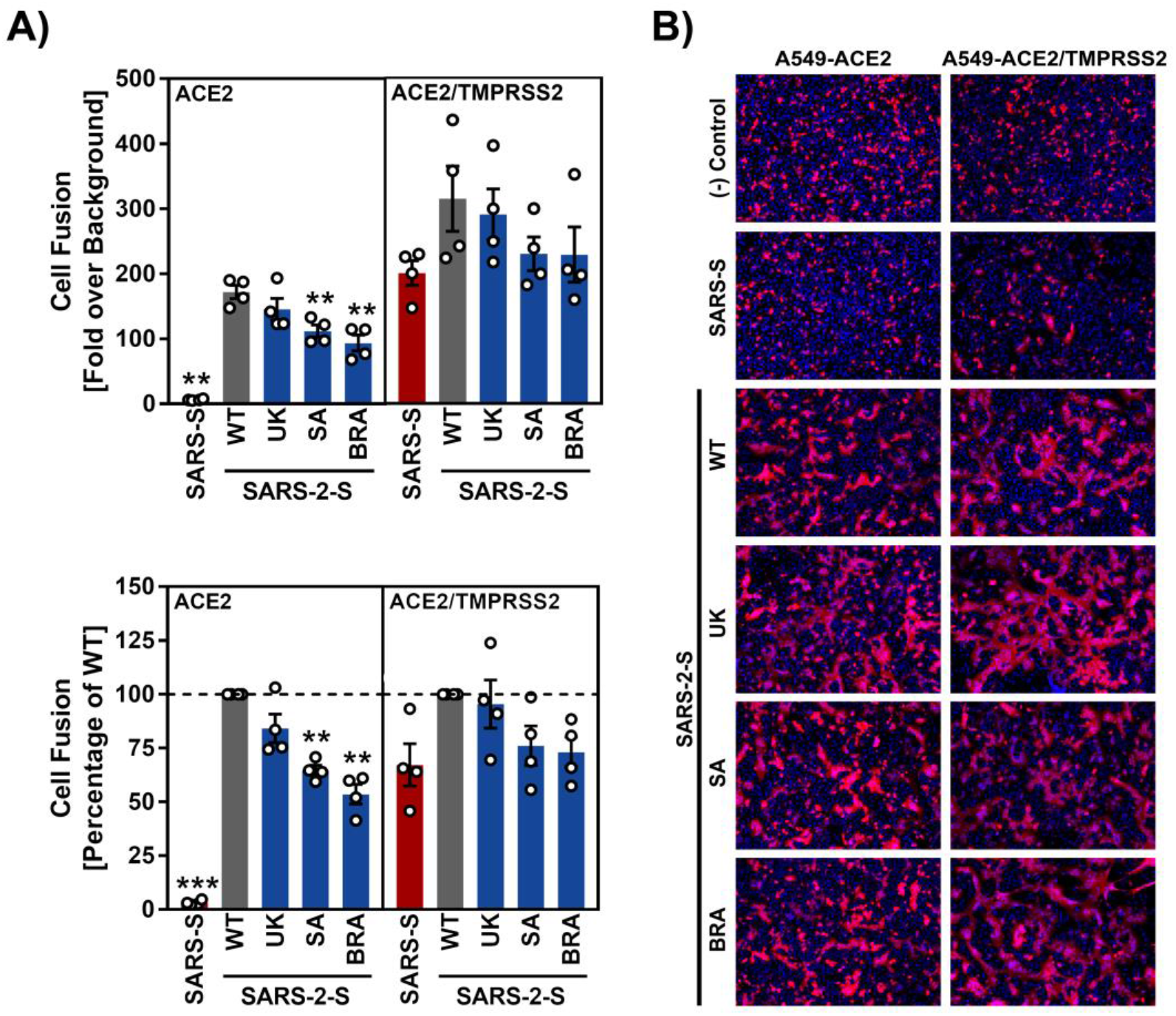
The S proteins of the SARS-CoV-2 variants drive robust cell-cell fusion. (A) Quantitative cell-cell fusion assay. S protein-expressing effector cells were mixed with ACE2 or ACE2/TMPRSS2 expressing target cells and cell-cell fusion was analyzed by measuring luciferase activity in cell lysates. Presented are the average (mean) data from four biological replicates. Error bars indicate the SEM. Statistical significance was analyzed by one-way ANOVA with Dunnett’s posttest. (B) Qualitative fusion assay. A549-ACE2 (left) and A549-ACE2/TMPRSS2 (right) cells were transfected to express the indicated S proteins (or no viral protein) along with DsRed. At 24 h posttransfection, cells were fixed and analyzed for the presence of syncytia by fluorescence microscopy (magnification: 10x). Presented are representative images from a single experiment. Data were confirmed in three additional experiments. WT = wildtype, GB = Great Britain, SA = South Africa, BRA = Brazil

### Similar stability and entry kinetics of particles bearing WT and variant S proteins

We next investigated whether the S proteins of the SARS-CoV-2 variants showed altered stability, which may contribute to the alleged increased transmissibility of the viral variants. For this, we incubated S protein-bearing particles for different time intervals at 33°C, a temperature that is present in the nasal cavity, and subsequently assessed their capacity to enter target cells. The efficiency of cell entry markedly decreased upon incubation of particles at 33°C for more than 8 h, but no appreciable differences were observed between particles bearing S proteins from SARS-CoV-2 WT or variants (Fig. 4A).

**Figure 4.**
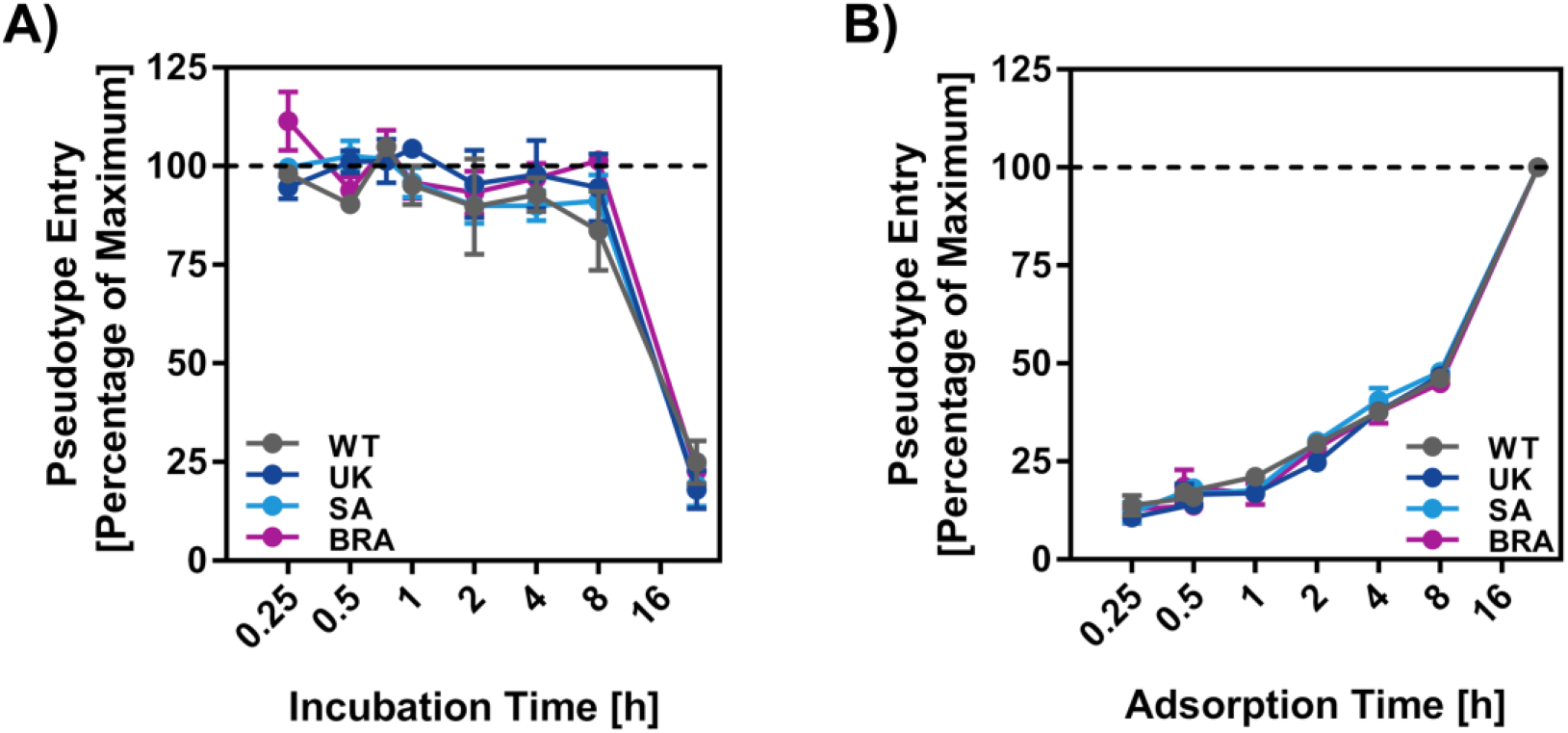
Particles bearing the S proteins of SARS-CoV-2 variants exhibit similar stability and entry kinetics. (A) Particles bearing the indicated S proteins were incubated for different time intervals at 33 °C, snap frozen, thawed and inoculated onto Vero cells. Entry of particles that were frozen immediately was set as 100%. (B) Particles bearing the indicated S proteins were incubated for indicated time intervals with Vero cells. Subsequently, the cells were washed and luciferase activity determined. Transduction measured without particle removal by washing was set as 100%. For both panels, the average (mean) data from three biological replicates (each performed with technical quadruplicates) is presented. Error bars indicate the SEM. Statistical significance was analyzed by one-way ANOVA with Dunnett’s posttest. WT = wildtype, GB = Great Britain, SA = South Africa, BRA = Brazil

Although the S proteins of the SARS-CoV-2 variants under study did not differ markedly from WT S protein regarding stability and entry efficiency, they might mediate entry with different kinetics as compared to WT S protein. To investigate this possibility, we incubated target cells with S protein-bearing particles for the indicated time intervals, removed unbound virus by washing and universally determined entry efficiency at 16 h post inoculation. Entry efficiency increased with the time available for particle adsorption to cells but no clear differences were observed between particles bearing WT S protein or S protein from SARS-CoV-2 variants (Fig. 4B). Our results suggest that there might be no major differences between WT SARS-CoV-2 and SARS-CoV-2 variants UK, South Africa and Brazil regarding S protein stability and entry kinetics.

### Soluble ACE2, TMPRSS2 inhibitors and membrane fusion inhibitors block entry

Soluble ACE2 (sACE2) blocks SARS-CoV-2 entry into cells and is in clinical development for COVID-19 therapy (Monteil et al., 2020). Similarly, the clinically proven protease inhibitors Camostat and Nafamostat block TMPRSS2-dependent SARS-CoV-2 cell entry and their potential for COVID-19 treatment is currently being assessed (Hoffmann et al., 2020b; Hoffmann et al., 2020c). Finally, the membrane fusion inhibitor EK1 and its optimized lipid-conjugated derivative EK1C4 block SARS-CoV-2 entry by preventing conformational rearrangements in S protein required for membrane fusion (Xia et al., 2020). We asked whether entry driven by the S proteins of UK, South Africa and Brazil variant can be blocked by these inhibitors. All inhibitors were found to be active although entry mediated by the S proteins of the SARS-CoV-2 variants was slightly more sensitive to blockade by sACE2 as compared to WT S protein, at least for certain sACE2 concentrations (Fig. 5). Conversely, entry driven by the S protein of the Brazil variant was slightly more sensitive to blockade by EK1 and EK1C4 as compared to the other S proteins tested (Fig. 5). These results suggest that sACE2, TMPRSS2 inhibitors and membrane fusion inhibitors will be active against UK, South Africa and Brazil variant.

**Figure 5.**
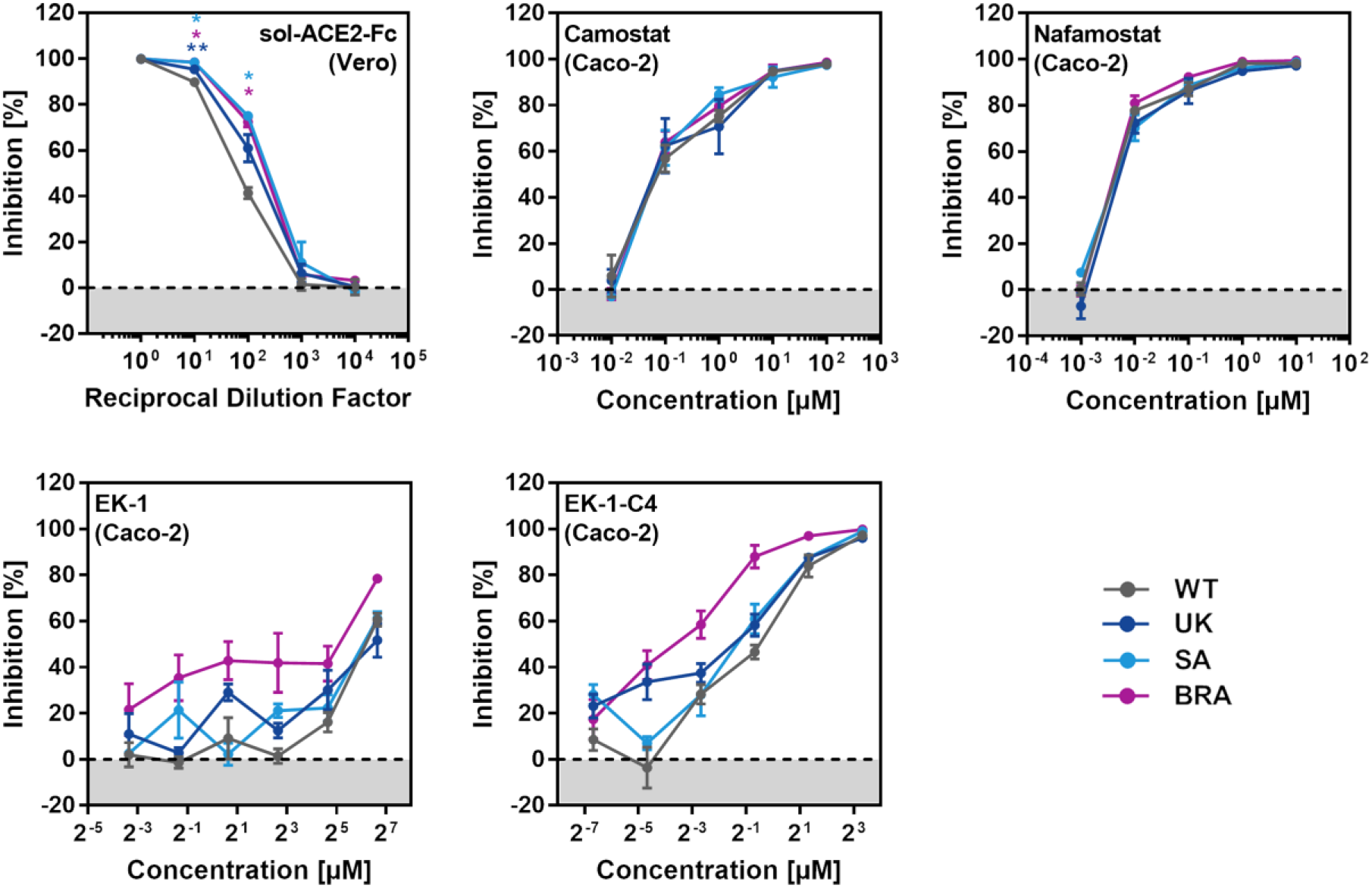
Entry driven by the S proteins of the SARS-CoV-2 variants can be blocked with soluble ACE2, protease inhibitors targeting TMPRSS2 and a membrane fusion inhibitor. Top row, left panel: S protein-bearing particles were incubated with different concentrations of soluble ACE2 (30 min, 37 °C) before being inoculated onto Vero cells. Top row, middle and right panel: Caco-2 target cells were pre-incubated with different concentrations of serine protease inhibitors (Camostat or Nafamostat; 1 h, 37 °C) before being inoculated with particles harboring the indicated S proteins. Bottom row, both panels: The peptidic fusion inhibitor EK-1 and its improved lipidated derivate EK-1-C4 were incubated with partices at indicated concentrations (30 min, 37 °C) and then added to Vero cells. All panels: Transduction efficiency was quantified by measuring virus-encoded luciferase activity in cell lysates at 16-20 h posttransduction. For normalization, inhibition of SARS-2-S-driven entry in samples without soluble ACE2 or inhibitor was set as 0 %. Presented are the average (mean) data from three biological replicates (each performed in technical triplicates [EK-1, EK-1-C4] or quadruplicates [soluble ACE2, Camostat, Nafamostat]). Error bars indicate the SEM. Statistical significance was analyzed by one-way ANOVA with Dunnett’s posttest. WT = wildtype, GB = Great Britain, SA = South Africa, BRA = Brazil

### Resistance against antibodies used for COVID-19 treatment

A cocktail of monoclonal antibodies (REGN-COV2, Casirivimab and Imdevimab) and the monoclonal antibody Bamlanivimab block SARS-CoV-2 WT infection and have received EUA for COVID-19 therapy. We analyzed whether these antibodies can inhibit entry driven by the S proteins of UK, South Africa and Brazil variants. All variants were comparably inhibited by antibody REGN10987 (Imdevimab) (Fig. 6). In contrast, entry driven by the S proteins of the South Africa and Brazil variant was partially resistant against antibody REGN10933 (Casirivimab) and fully resistant against REGN10989 (Fig. 6). Finally, entry mediated by the S proteins of the South Africa and Brazil variant was completely resistant to Bamlanivimab while the S protein of the UK variant was efficiently blocked by all antibodies tested (Fig. 6). Collectively, these data indicate that antibodies with EUA might provide incomplete (REGENERON) or no (Bamlanivimab) protection against the South Africa and Brazil variants.

**Figure 6.**
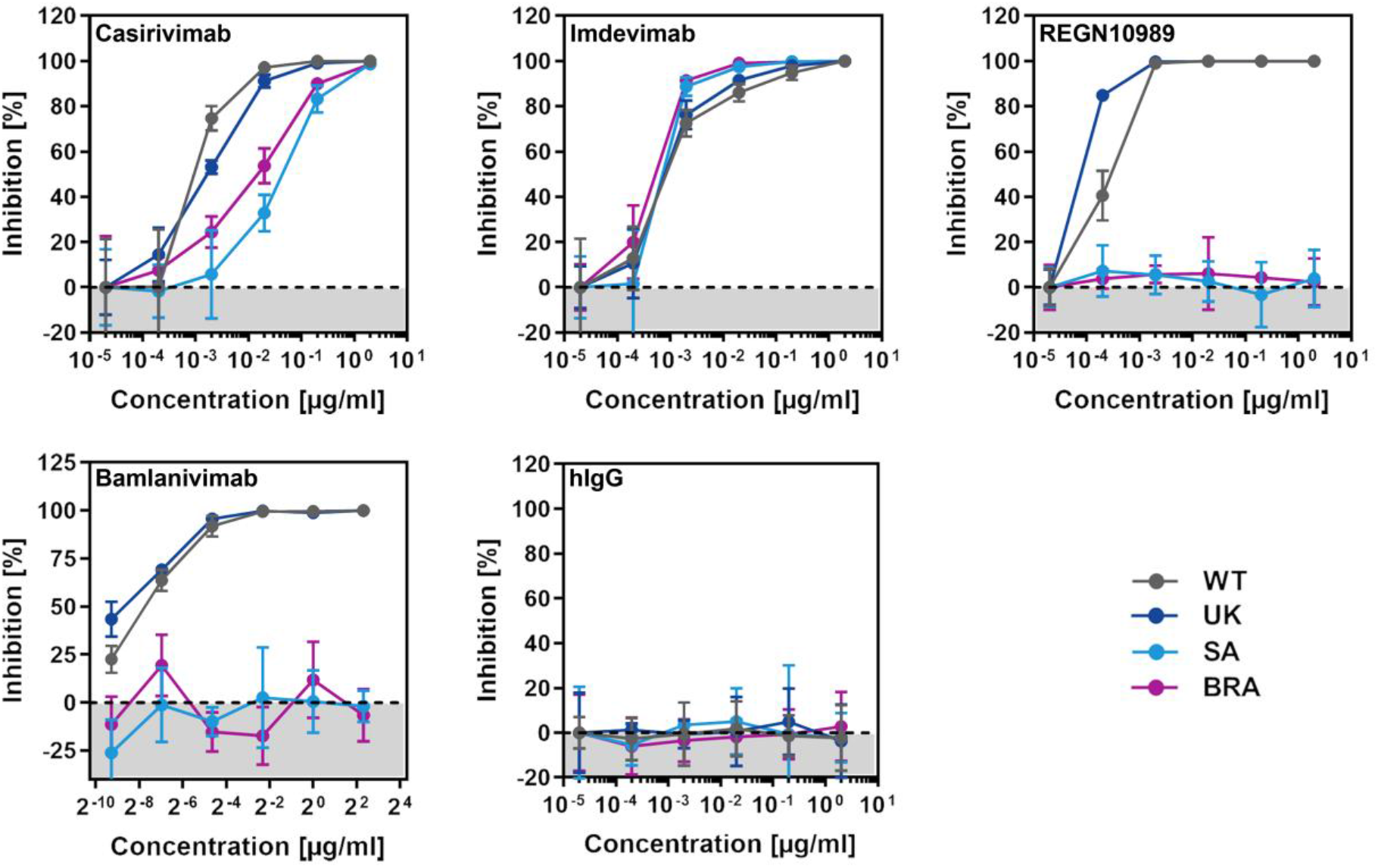
The S proteins of SARS-CoV-2 variants from South Africa and Brazil are partially or fully resistant to inhibition by therapeutic monoclonal antibodies with EUA. Pseudotypes bearing the indicated S proteins were incubated (30 min, 37 °C) with different concentrations of control antibody (hIgG), three different Regeneron antibodies (Casirivimab, Imdevimab, REGN10989) or Bamlanivimab (LY-CoV555) before being inoculated onto target Vero cells. Transduction efficiency was quantified by measuring virus-encoded luciferase activity in cell lysates at 16-20 h posttransduction. For normalization, inhibition of S protein-driven entry in samples without antibody was set as 0 %. Presented are the data from a single experiment performed with technical triplicates. Data were confirmed in a separate experiment. Error bars indicate standard deviation (SD). WT = wildtype, GB = Great Britain, SA = South Africa, BRA = Brazil

### Reduced neutralization by plasma from convalescent patients

SARS-CoV-2 infection can induce the production of neutralizing antibodies and these antibodies are believed to contribute to protection from reinfection (Rodda et al., 2020; Wajnberg et al., 2020). Therefore, it is important to elucidate whether UK, South Africa and Brazil variants are efficiently neutralized by antibody responses in convalescent COVID-19 patients. We addressed this question using plasma collected from COVID-19 patients undergoing intensive care at Göttingen University Hospital, Germany. The plasma samples had been pre-screened for high neutralizing activity against WT S protein, and a plasma sample with no neutralizing activity was included as negative control. Spread of SARS-CoV-2 variants in Germany was very limited at the time of sample collection, indicating that serum antibodies were induced in response to SARS-CoV-2 WT infection.

All plasma samples with known neutralizing activity (ID15, 18, 20, 22, 23, 24, 27, 33, 51) efficiently reduced entry driven by WT S protein while the control plasma (ID16) was inactive (Fig. 7A). Blockade of entry driven by the S protein of the UK variant was slightly less efficient (Fig. 7A and C). In contrast, seven out of nine plasma samples inhibited entry driven by the S proteins of the South Africa and Brazil variants less efficiently as compared to entry driven by WT S protein. These results suggest that individuals previously infected with WT SARS-CoV-2 might only be partially protected against infection with South Africa and Brazil variants of SARS-CoV-2.

**Figure 7.**
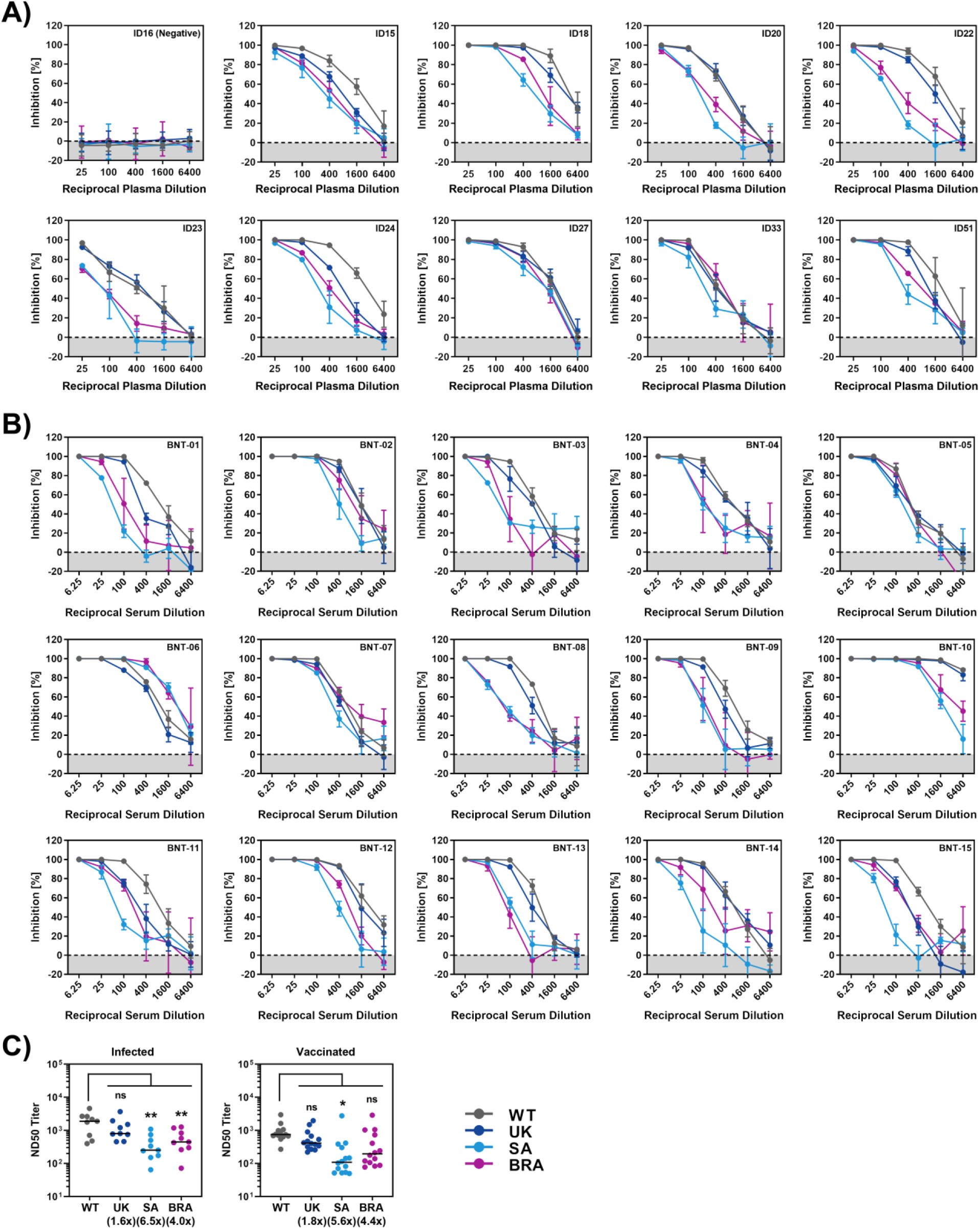
S proteins of SARS-CoV-2 variants from South Africa and Brazil show reduced neutralization sensitivity against convalsecent plasma and serum from BNT162b2 vaccinated individuals. Pseudotypes bearing the indicated S proteins were incubated (30 min, 37 °C) with different dilutions of plasma derived from COVID-19 patients (A) or serum from individuals vaccinated with the Pfizer/BioNTech vaccine BNT162b2 (obtained 13-15 days after the second dose) and inoculated onto Vero target cells. Transduction efficiency was quantified by measuring virus-encoded luciferase activity in cell lysates at 16-20 h posttransduction. The results are shown as % inhibition. For normalization, S protein-driven entry in the absence of plasma/serum was set as 0 %. Presented are the data from a single experiment performed with technical triplicates. Error bars indicate SD. Results were confirmed in a second biological replicate. (C) Serum dilutions that lead to a 50% reduction in S protein-driven transduction (neutralization titer 50, NT50) were calculated for convalsecent plasma (left) and vaccinee sera (right). Presented are the data derived from panels A and B. The line indicates the median. WT = wildtype, GB = Great Britain, SA = South Africa, BRA = Brazil

### Reduced neutralization by sera from BNT162b2-vaccinated individuals

The vaccine BNT162b2 is based on an mRNA that encodes for the viral S protein and is highly protective against COVID-19 (Polack et al., 2020). While the S protein harbor T-cell epitopes (Grifoni et al., 2020; Peng et al., 2020), efficient protection is believed to require the induction of neutralizing antibodies. We determined neutralizing activity of sera from 15 donors immunized twice with BNT162b2 (Table S1). All sera efficiently inhibited entry driven by the WT S protein and inhibition of entry driven by the S protein of the UK variant was only slightly reduced (Fig. 7B,C). In contrast, 12 out of 15 sera showed a markedly reduced inhibition of entry driven by the S proteins of the South Africa and Brazil variant (Fig. 7B,C), although it should be stated that all sera completely inhibited entry at the lowest dilution tested. In sum, these results suggest that BNT162b2 may offer less robust protection against infection by these variants as compared to SARS-CoV-2 WT.

## DISCUSSION

The COVID-19 pandemic has taken a major toll on human health and prosperity. Non-pharmaceutic interventions are currently the major instrument to combat the pandemic but are associated with a heavy burden on economies. Protective vaccines became recently available and might become a game changer – it is hoped that efficient vaccine roll out might allow to attain herd immunity in certain countries in the second half of 2021. The recent emergence of SARS-CoV-2 variants UK, South Africa and Brazil that harbor mutations in the major target of neutralizing antibodies, the viral S protein, raises the question whether vaccines available at present will protect against infection with these viruses. Similarly, it is largely unclear whether antibody responses in convalescent patients protect against re-infection with the new variants. The results of the present study suggest that SARS-CoV-2 variants South Africa and Brazil are partially (Casirivimab) or fully (Bamlanivimab) resistant against antibodies used for COVID-19 treatment and are inhibited less efficiently by convalescent plasma or sera from individuals immunized with the mRNA vaccine BNT162b2. These results suggest that strategies relying on antibody-mediated control of SARS-CoV-2 infection might be compromised by resistance development.

The increased transmissibility postulated for the UK variant and purported for the South Africa and Brazil variants suggest that these viruses might exhibit altered host-cell interactions or stability. The present analysis suggests that there are no major differences in host cell entry of WT SARS-CoV-2 and the UK, South Africa and Brazil variant (CDC, 2021; Leung et al., 2021). Thus, the S proteins of these viruses mediated entry into various cell lines with roughly comparable efficiency and no evidence for increased S protein stability or differences in entry kinetics were obtained. Similarly, the S proteins of all variants were able to mediate fusion of human cells. Moreover, entry driven by all S proteins studied was blocked by sACE2, protease inhibitors targeting TMPRSS2 and a membrane fusion inhibitor. However, it should be noted that the S proteins of all variants were slightly more susceptible to blockade by sACE2, suggesting differences in ACE2 engagement between WT and variant S proteins.

Although host-cell interactions underlying viral entry might not differ markedly between SARS-CoV-2 S protein WT and the variants studied here, major differences in susceptibility to antibody-mediated neutralization were observed. Entry driven by the S proteins of the South Africa and Brazil variants was not inhibited by one of the REGENERON antibodies (REGN10989) and Bamlanivimab (Baum et al., 2020a; Baum et al., 2020b; Chen et al., 2020; Gottlieb et al., 2021), suggesting that these antibodies might not be suitable for treatment of COVID-19 patients infected with these variants. The partial resistance against Casirivimab (REGN10933) is in keeping with mutations present in the S protein of South Africa and Brazil variant being located at the antibody binding site (Fig. S1). Moreover, and more importantly, entry driven by the S proteins of the South Africa and Brazil variants were markedly less sensitive to neutralization by antibodies from convalescent patients and vaccinated individuals as compared to the WT S protein. It should be noted that all plasma and sera tested completely inhibited entry at the lowest dilution tested and that T cell responses will contribute to control of SARS-CoV-2 infection, particularly in re-infected convalescent patients (Grifoni et al., 2020; Peng et al., 2020). Nevertheless, the markedly reduced sensitivity to antibody-mediated neutralization suggests that convalescent and vaccinated individuals might not be fully protected against infection by the South Africa and Brazil variants. Such a scenario would be in keeping with preliminary information suggesting that certain vaccines might provide less effective protection in South African as compared to the US (Callaway and Mallapaty, 2021). On a more general level, our findings suggest that the interface between the SARS-CoV-2 S protein and ACE2 exhibits high plasticity, favoring emergence of escape variants.

Our find that entry driven by the S protein of the UK variant can be efficiently inhibited by antibodies induced upon infection and vaccination is in agreement with those of Muik and colleagues, who reported that pseudoparticles bearing the UK S protein are efficiently neutralized by sera from BNT162b2 vaccinated individuals (Muik et al., 2021). Xie and coworkers found that authentic SARS-CoV-2 bearing two mutations present in the S protein of the UK variant (69/70-deletion + N501Y) was still robustly neutralized by antibodies induced by vaccination with BNT162b2, again in keeping with our findings. Neutralization of a virus bearing two changes found in the S protein of the South Africa variant (E484K + N501Y) was moderately reduced and it is conceivable that neutralization resistance would have been further increased by the other four mutations present in the S1 unit of the S protein of the South Africa variant, including K417N, which is located in the RBD (Xie et al., 2021).

Our results await confirmation with authentic SARS-CoV-2. However, the data available at present suggest that the South Africa and Brazil variants constitute an elevated threat to human health and that containment of these variants by non-pharmaceutic interventions is an important task.

## MATERIAL AND METHODS

### Cell culture

All cell lines were incubated at 37 °C in a humidified atmosphere containing 5% CO2. All media were supplemented with 10% fetal bovine serum (FCS, Biochrom or GIBCO), 100 U/ml of penicillin and 0.1 mg/ml of streptomycin (PAN-Biotech). 293T (human, kidney; ACC-635, DSMZ), 293T cells stably expressing ACE2 (293T-ACE2), BHK-21 (Syrian hamster, kidney cells; CCL-10, ATCC), Vero76 (African green monkey, kidney; CRL-1586, ATCC; kindly provided by Andrea Maisner, Institute of Virology, Philipps University Marburg, Marburg, Germany) and Vero-TMPRSS2 cells (Hoffmann et al., 2020b) were cultivated in Dulbecco’s modified Eagle medium (DMEM). Vero-TMPRSS2 cells additionally received puromycin (0.5 μg/ml, Invivogen). A549 (human, lung; CRM-CCL-185, ATCC), A549-ACE2 and A549-ACE2/TMPRSS2 cells were cultivated in DMEM/F-12 Medium with Nutrient Mix (ThermoFisher Scientific). A549-ACE2 cells further received 0.5 μg/ml puromycin, while A549-ACE2/TMPRSS2 cells were cultivated in the presence of 0.5 μg/ml puromycin and 1 μg/ml blasticidin. Caco-2 (human, intestine; HTB-37, ATCC) and Calu-3 cells (human, lung; HTB-55, ATCC; kindly provided by Stephan Ludwig, Institute of Virology, University of Münster, Germany) were cultivated in minimum essential medium supplemented with 1x non-essential amino acid solution (from 100x stock, PAA) and 1 mM sodium pyruvate (Thermo Fisher Scientific). 293T cells that stably express ACE2 were generated by retroviral (murine leukemia virus, MLV) transduction and selection of parental 293T cells with puromycin (4 μg/ml for initial selection and 0.5 μg/ml for sub-culturing). Similarly, we generated A549 cells stably expressing ACE2 (A549-ACE2). A549 cells stably expressing ACE2 and TMPRSS2 (A549-ACE2/TMPRSS2) were obtained by retroviral transduction of A549-ACE2 cells and selection with blasticidin (6 μg/ml for initial selection and 1 μg/ml for sub-culturing). Authentication of cell lines was performed by STR-typing, amplification and sequencing of a fragment of the cytochrome c oxidase gene, microscopic examination and/or according to their growth characteristics. Further, cell lines were routinely tested for contamination by mycoplasma. Transfection of cells was carried out by the calcium-phosphate method or by using polyethylenimin, Lipofectamine LTX (Thermo Fisher Scientific) or Transit LT-1 (Mirus).

### Plasmids

Expression plasmids for DsRed (PMID: 32142651), vesicular stomatitis virus (VSV, serotype Indiana) glycoprotein (VSV-G) (Brinkmann et al., 2017), SARS-S (derived from the Frankfurt-1 isolate; containing a C-terminal HA epitope tag) (Hoffmann et al., 2020b), SARS-2-S (codon-optimized, based on the Wuhan/Hu-1/2019 isolate; with a C-terminal truncation of 18 amino acid residues or with a C-terminal HA epitope tag) (Hoffmann et al., 2020b), angiotensin-converting enzyme 2 (ACE2) (Hoffmann et al., 2013), TMPRSS2 (Heurich et al., 2014) have been described elsewhere. In order to generate expression vectors for S proteins from emerging SARS-CoV-2 variants, we introduced the required mutations into the parental SARS-2-S sequence by overlap extension PCR. Subsequently, the respective open reading frames were inserted into the pCG1 plasmid (kindly provided by Roberto Cattaneo, Mayo Clinic College of Medicine, Rochester, MN, USA), making use of the unique BamHI and XbaI restriction sites. Further, we cloned the coding sequence for human ACE2 into the pQCXIP plasmid (Brass et al., 2009), yielding pQCXIP_ACE2. For the generation of cell lines stably overexpressing human TMPRSS2 and/or human ACE2 we produced MLV-based transduction vectors using the pQCXIBl_cMYC-hTMPRSS2 (Kleine-Weber et al., 2018) or pQCXIP_ACE2 expression vectors in combination with plasmids coding for VSV-G and MLV-Gag/Pol (Bartosch et al., 2003). In order to obtain the expression vector for soluble human ACE2 harboring the Fc portion of human immunoglobulin G (sol-ACE2-Fc), we PCR amplified the sequence coding for the ACE2 ectodomain (amino acid residues 1-733) and cloned it into the pCG1-Fc plasmid ((Sauer et al., 2014), kindly provided by Georg Herrler, University of Veterinary Medicine, Hannover, Germany). Sequence integrity was verified by sequencing using a commercial sequencing service (Microsynth Seqlab). Specific cloning details (e.g., primer sequences and restriction sites) are available upon request.

### Sequence analysis and protein models

S protein sequences of emerging SARS-CoV-2 S variants found in the United Kingdom (UK, EPI_ISL_601443), South Africa (SA, EPI_ISL_700428) and Brazil (BRA, EPI_ISL_792683) were retrieved from the GISAID (global initiative on sharing all influenza data) database (https://www.gisaid.org/). Protein models are based on PDB: 6XDG (Hansen et al., 2020) or a template generated by modelling the SARS-2-S sequence on a published crystal structure (PDB: 6XR8,(Cai et al., 2020)), using the SWISS-MODEL online tool (https://swissmodel.expasy.org/), and were generated using the YASARA software (http://www.yasara.org/index.html).

### Production of soluble ACE2 (sol-ACE2-Fc)

293T cells were grown in a T-75 flask and transfected with 20 μg of sol-ACE2-Fc expression plasmid. At 10 h posttransfection, the medium was replaced and cells were further incubated for 38 h before the culture supernatant was collected and centrifuged (2,000 x g, 10 min, 4 °C). Next, the clarified supernatant was loaded onto Vivaspin protein concentrator columns with a molecular weight cut-off of 30 kDa (Sartorius) and centrifuged at 4,000 x g, 4 °C until the sample was concentrated by a factor of 20. The concentrated sol-ACE2-Fc was aliquoted and stored at −80 °C until further use.

### Collection of serum and plasma samples

Sera from individuals vaccinated with BioNTech/Pfizer vaccine BNT162b2 were obtained 13-15 days after the second dose. Study was approved by the Ethic committee of Ulm university (vote 31/21 – FSt/Sta). Collection of plasma samples from COVID-19 patients treated at the intensive care unit was approved by the Ethic committee of the University Medicine Göttingen (SeptImmun Study 25/4/19 Ü). For collection of plasma, Cell Preparation Tube (CPT) vacutainers with sodium citrate were used and plasma was collected as supernatant over the PBMC layer. For vaccinated patients, blood was collected in S-Monovette^®^ Serum Gel tubes (Sarstedt). Subsequently, the plasma and serum samples were incubated at 56°C for 30 min to inactivate putative infectious virus and for reconvalescent plasma pre-screening for detection of neutralizing activity was performed on Vero76 cells using SARS-2-S- and VSV-G bearing pseudotypes as control, normalized for equal infectivity.

### Pseudotyping of VSV and transduction experiments

Rhabdoviral pseudotype particles were prepared according to a published protocol (Kleine-Weber et al., 2019). For pseudotyping we used a replication-deficient VSV vector that lacks the genetic information for VSV-G and instead codes for two reporter proteins, enhanced green fluorescent protein and firefly luciferase (FLuc), VSV*ΔG-FLuc (kindly provided by Gert Zimmer, Institute of Virology and Immunology, Mittelhäusern, Switzerland) (Berger Rentsch and Zimmer, 2011). 293T cells transfected to express the desired viral glycoprotein were inoculated with VSV*ΔG-FLuc and incubated for 1 h at 37 °C before the inoculum was removed and cells were washed. Finally, culture medium containing anti-VSV-G antibody (culture supernatant from I1-hybridoma cells; ATCC no. CRL-2700) was added. Following an incubation period of 16-18 h, pseudotype particles were harvested by collecting the culture supernatant, pelleting cellular debris through centrifugation (2,000 x g, 10 min, room temperature) and transferring aliquots of the clarified supernatant into fresh reaction tubes. Samples were stored at −80 °C. For transduction experiments, target cells were seeded in 96-well plates, inoculated with the respective pseudotype particles with comparable infectivity and further incubated. At 16-18 h postinoculation, transduction efficiency was analyzed. For this, the culture supernatant was removed and cells were lysed by incubation for 30 min at room temperature with Cell Culture Lysis Reagent (Promega). Next, lysates were transferred into white 96-well plates and FLuc activity was measured using a commercial substrate (Beetle-Juice, PJK; Luciferase Assay System, Promega) and a plate luminometer (Hidex Sense Plate Reader, Hidex or Orion II Microplate Luminometer, Berthold)..

Depending on the experimental set-up target cells were either transfected in advance (24 h) with ACE2 expression plasmid or empty vector (BHK-21), or pre-incubated with different concentrations of serine protease inhibitor (Camostat or Nafamostat, Caco-2, 1 h at 37 °C). Alternatively, pseudotype particles were pre-incubated with different concentrations of either sol-ACE2-Fc, fusion inhibitor (EK-1 or EK-1-C4), monoclonal antibodies (REGN10933, REGN10987, REGN10989, Bamlaivimab/LY-CoV555), or sera from COVID-19 patients or vaccinated (Pfizer/BioNTech vaccine BNT162b2) individuals (30 min at 37 °C). S protein stability was analyzed as follows, pseudotype particles were incubated for different time intervals at 33 °C the snap-frozen and stored at −80 °C until all samples were collected. Thereafter, samples were thawed and inoculated onto Vero76 cells and incubated as described above. For the investigation of the entry speed of S protein-bearing pseudotypes, the respective particles were inoculated on Vero76 cells and adsorbed for different time intervals before the inoculum was removed and cells were washed and incubated with fresh medium.

### Analysis of S protein expression by fluorescence microscopy

A549-ACE2 cells that were grown on coverslips were transfected with plasmids encoding SARS-CoV-2 S protein variants with a C-terminal HA epitope tag or empty expression vector (control). At 24 h posttransfection, cells were fixed with 4 % paraformaldehyde solution (30 min, room temperature), washed and incubated (15 min, room temperature) with phosphate-buffered saline (PBS) containing 0.1 M glycine and permeabilized by treatment with 0.2 % Triton-X-100 solution (in PBS, 15 min). Thereafter, samples were washed and incubated for 1 h at room temperature with primary antibody (anti-HA, mouse, 1:500, Sigma-Aldrich) diluted in PBS containing 1 % bovine serum albumin. Next, the samples were washed with PBS and incubated in the dark for 1 h at 4 °C with secondary antibody (Alexa Fluor-568-conjugated anti-mouse antibody, 1:750, Thermo Fisher Scientific). Finally, the samples were washed, nuclei were stained with DAPI and coverslips were mounted onto microscopic glass slides with Mowiol/DABCO. Images were taken using a Zeiss LSM800 confocal laser scanning microscope with ZEN imaging software (Zeiss).

### Qualitative cell-cell fusion assay

A549-ACE2 or A549-ACE2/TMPRSS2 cells were transfected with DsRed expression plasmid along with either expression vector for wildtype or mutant SARS-2-S, SARS-S or empty plasmid. At 24 h posttransfection, cells were fixed with 4 % paraformaldehyde solution (30 min, room temperature), washed and nuclei were stained with DAPI. Next, cells were washed again with PBS and images were taken using a Zeiss LSM800 confocal laser scanning microscope with ZEN imaging software (Zeiss).

### Quantitative cell-cell fusion assay

293T target-cells were seeded in a 48-well plate at 50.000 cells/well and transfected with Gal4-TurboGFP-Luciferase expression plasmid (Gal4-TurboGFP-Luc) as well as expression plasmid for ACE2 alone or in combination with TMPRSS2 (5:1 ratio). 293T effector-cells were seeded in a 10 cm dish at 70-80% confluency and transfected with the Vp16-Gal4 expression plasmid as well as expression plasmid for WT or mutant SARS-2-S, SARS-S or empty plasmid. At 24h posttransfection, effector-cells were detached by resuspending them in culture medium and added to the target-cells in a 1:1 ratio. After 24-48 h luciferase activity was analyzed using the PromoKine Firefly Luciferase Kit or Beetle-Juice Luciferase Assay according to manufacturer’s instructions and a Biotek Synergy 2 plate reader.

### Data normalization and statistical analysis

Data analysis was performed using Microsoft Excel as part of the Microsoft Office software package (version 2019, Microsoft Corporation) and GraphPad Prism 8 version 8.4.3 (GraphPad Software). Data normalization was done as follows: (i) To compare efficiency of cell entry driven by the different S protein variants under study, transduction was normalized against SARS-CoV-2 S WT (set as 100%); (ii) For experiments investigating inhibitory effects, transduction was normalized against a reference sample (e.g., control-treated cells or pseudotypes, set as 100%). Serum dilutions the cause a 50 % reduction of transduction efficiency (neutralizing titer 50, NT50), were calculated using a non-linear regression model (inhibitor vs. normalized response, variable slope). Statistical significance was tested by one- or two-way analysis of variance (ANOVA) with Dunnett’s or Sidak’s post-hoc test, or by paired student’s t-test. Only P values of 0.05 or lower were considered statistically significant (P > 0.05, not significant [ns]; P ≤ 0.05, *; P ≤ 0.01, **; P ≤ 0.001, ***). Specific details on the statistical test and the error bars are indicated in the figure legends.

## ACKNOWLEDGMENTS

J.M. acknowledges funding by a Collaborative Research Centre grant of the German Research Foundation (316249678 – SFB 1279), the European Union’s Horizon 2020 research and innovation programme under grant agreement No 101003555 (Fight-nCoV) and the Federal Ministry of Economics, Germany (Combi-CoV-2). J.M. and A.K. acknowledge funding by the Ministry for Science, Research and the Arts of Baden-Württemberg, Germany, and the German Research Foundation (Fokus-Förderung COVID-19). R.G. A.S. and R.GA.S.. are part of the International Graduate School in Molecular Medicine Ulm. S.P. acknowledges funding by BMBF (RAPID Consortium, 01KI1723D and 01KI2006D; RENACO, 01KI20328A, 01KI20396, COVIM consortium), the county of Lower Saxony and the German Research Foundation (DFG). H.S. acknowledges funding from the German Federal Ministry of Health. H.S. acknowledges funding from the Ministry for Science, Research and the Arts of Baden-Württemberg, Germany; the European Commission (HORIZON2020 Project SUPPORT-E, no. 101015756) and the German Federal Ministry of Health. A.S.H. acknowledges funding from the German Research Foundation (HA 6013/6-1).

## AUTHOR CONTRIBUTIONS

Conceptualization, M.H., J.M., S.P.; Funding acquisition, S.P., J.M.; Investigation, M.H., P.A., R.G., A.S., B.H., A.H., N.K., L.G., H.H.-W., A.K., Essential resources, M.S.W., S.S., H.-M.J., B.J., H.S., M.M., A.K.; Writing, M.H. and S.P., Review and editing, all authors.

---

**Table S1:**
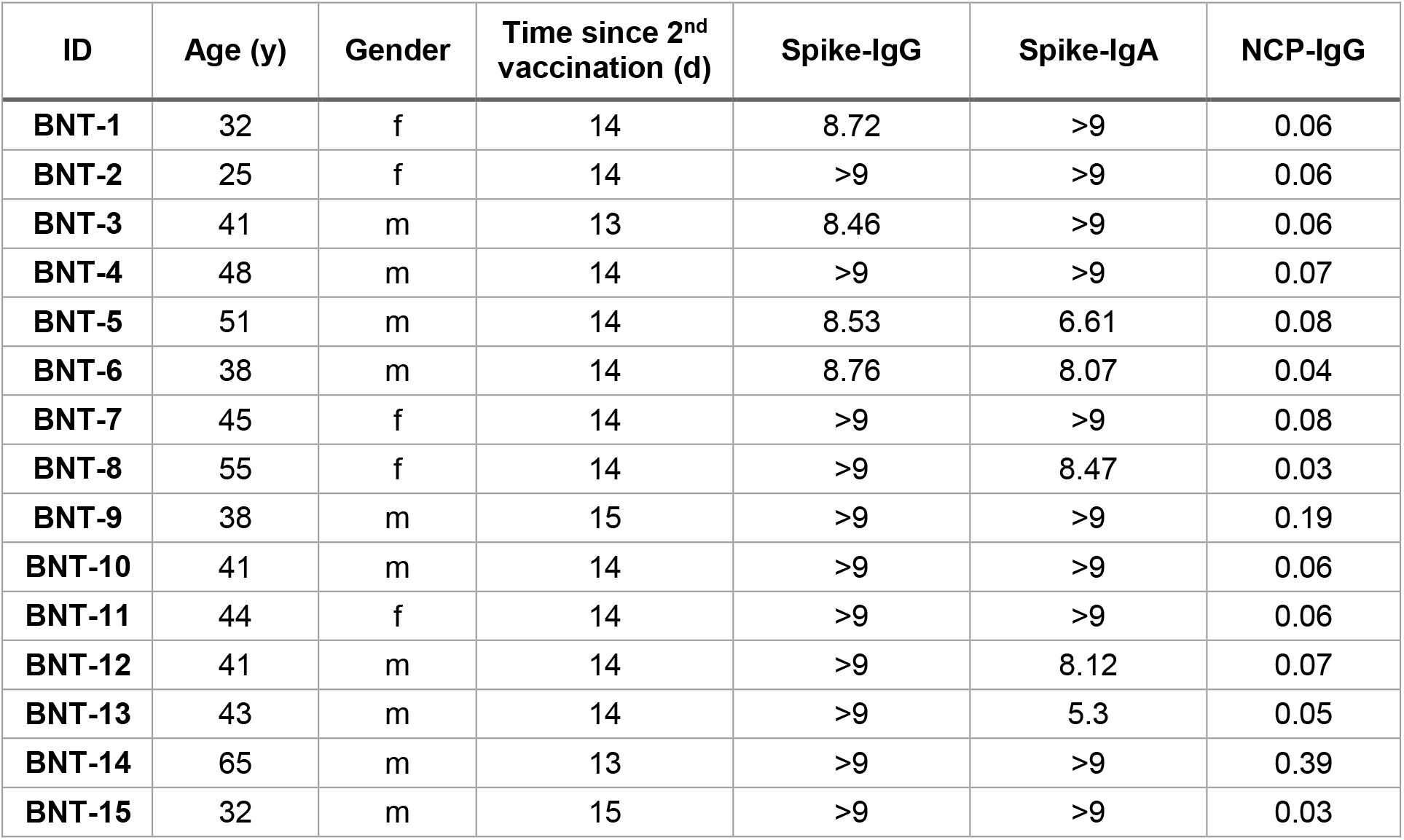
BNT162b2-vaccinated patient data. Serological data shows antibody titer against Spike (IgG, IgA) and Nucleocapsid (NCP, IgG) protein measured by Euroimmun-ELISA, values are given as baseline-corrected OD ratios compared to a calibrator. For all analytes, a ratio < 0.8 was considered to be non-reactive or negative. An OD-ratio of ≥ 1.1 was considered to be positive for all three analytes.

| ID | Age (y) | Gender | Time since 2<sup>nd</sup> vaccination (d) | Spike-IgG | Spike-IgA | NCP-IgG |
| --- | --- | --- | --- | --- | --- | --- |
| BNT-1 | 32 | f | 14 | 8.72 | >9 | 0.06 |
| BNT-2 | 25 | f | 14 | >9 | >9 | 0.06 |
| BNT-3 | 41 | m | 13 | 8.46 | >9 | 0.06 |
| BNT-4 | 48 | m | 14 | >9 | >9 | 0.07 |
| BNT-5 | 51 | m | 14 | 8.53 | 6.61 | 0.08 |
| BNT-6 | 38 | m | 14 | 8.76 | 8.07 | 0.04 |
| BNT-7 | 45 | f | 14 | >9 | >9 | 0.08 |
| BNT-8 | 55 | f | 14 | >9 | 8.47 | 0.03 |
| BNT-9 | 38 | m | 15 | >9 | >9 | 0.19 |
| BNT-10 | 41 | m | 14 | >9 | >9 | 0.06 |
| BNT-11 | 44 | f | 14 | >9 | >9 | 0.06 |
| BNT-12 | 41 | m | 14 | >9 | 8.12 | 0.07 |
| BNT-13 | 43 | m | 14 | >9 | 5.3 | 0.05 |
| BNT-14 | 65 | m | 13 | >9 | >9 | 0.39 |
| BNT-15 | 32 | m | 15 | >9 | >9 | 0.03 |

## SUPPLEMENTAL INFORMATION

**Fígure S1.**
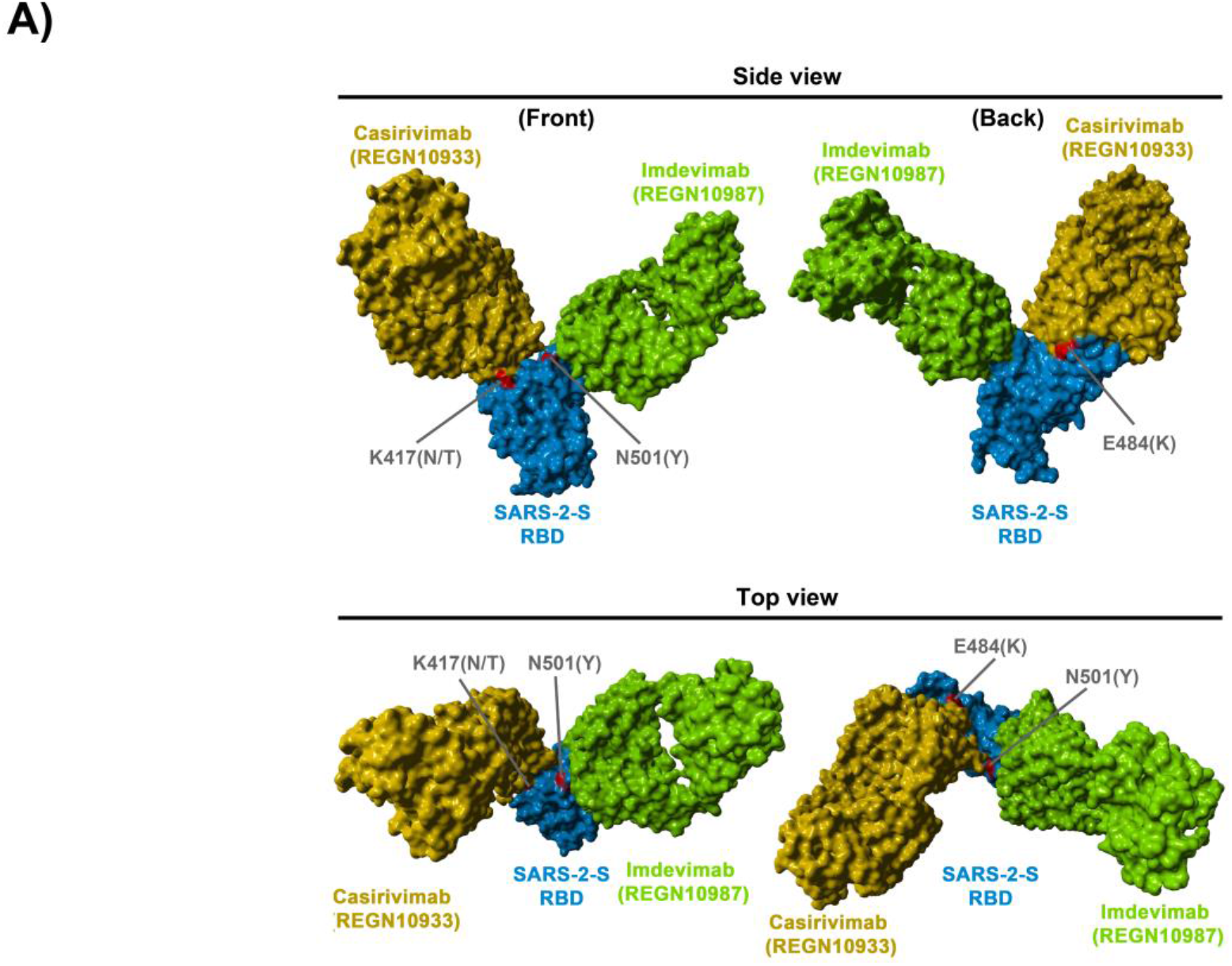
Location of SARS-2-S RBD mutations K417N/T, E484K and N501Y with respect to the binding interface of the REGN-COV2 antibody cocktail (related to Figure 6). The protein models of the SARS-2-S receptor-binding domain (RBD, blue) in complex with antibodies Casirivimab (REGN10933, orange) and Imdevimab (REGN10987, green) were constructed based on the 6XDG template (Hansen et al., 2020). Residues highlighted in red indicate amino acid variations found in emerging SARS-CoV-2 variants from the United Kingdom, South Africa and Brazil.

